# nf-core/proteinfamilies: A scalable pipeline for the generation of protein families

**DOI:** 10.1101/2025.08.12.670010

**Authors:** Evangelos Karatzas, Martin Beracochea, Fotis A. Baltoumas, Eleni Aplakidou, Lorna Richardson, James A. Fellows Yates, Daniel Lundin, nf-core community, Aydin Buluç, Nikos C. Kyrpides, Ilias Georgakopoulos-Soares, Georgios A. Pavlopoulos, Robert D. Finn

## Abstract

The growth of metagenomics-derived amino acid sequence data has transformed our understanding of protein function, microbial diversity and evolutionary relationships. However, the vast majority of these proteins remain functionally uncharacterized. Grouping the millions of such uncharacterised sequences with the few experimentally characterised ones allows the transfer of annotations, while the inspection of conserved residues with multiple sequence alignments can provide clues to function, even in the absence of existing functional information. To address the challenges associated with this data surge and the need to group sequences, we present a scalable, open-source, parametrizable Nextflow pipeline (*nf-core/proteinfamilies*) that generates protein nascent families or assigns new proteins to existing families. The computational benchmarks demonstrated that resource usage can scale approximately linearly with input size, while the biological benchmarks showed that the generated protein families closely resemble manually curated families found in widely used databases.

## Introduction

The generation of protein families is a common approach for facilitating the transfer of functional annotations from sequences with a determined function to uncharacterized sequences. Given the large volume of unannotated sequences that are obtained through metagenomic analyses, it is necessary to group sequences, in order to transfer such functional annotations at scale [1,2]. Grouping a few related sequences and building protein family models, can enable the detection and annotation of distant homologs across massive datasets. Profile Hidden Markov Models (HMMs) are especially effective for this task, as they capture conserved sequence patterns and model insertions and deletions. Efficiently transferring annotations at scale can expedite key applications such as identifying novel enzymes for industrial use, predicting drug targets in pathogenic organisms, and reconstructing metabolic pathways in microbial communities. To achieve this, biological databases and platforms provide curated protein information and metadata to support both functional and 3D-structure annotation of protein families. Major resources include UniProtKB [3], Protein Data Bank (PDB) [4], RefSeq [5], GenBank [6], IMG/M [7], Big Fantastic Database (BFD) [8], and MGnify [9]. These databases collectively host hundreds of millions to billions of sequences, many of which remain uncharacterized, particularly those derived from metagenomic sources.

While the aforementioned databases catalog raw sequence and structure data, protein classification resources help interpret this information by organizing proteins into families based on shared evolutionary or functional traits. Some widely used resources include Pfam [10], InterPro [11], the Novel Metagenome Protein Families Database (NMPFamsDB) [12,13], FunFams [14], eggNOG [15,16], KEGG Orthology (KO) [17], Clusters of Orthologous Groups (COG) [18] and its extension for eukaryotic proteins, KOG [19], each serving distinct purposes in protein classification. Notably InterPro integrates protein family, domain, and functional site data from thirteen contributing databases. As of May 2025, InterPro provides standardized annotations for over 200 million proteins. Conversely, NMPFamsDB focuses on protein families derived from metagenomic and metatranscriptomic data that do not correspond to known reference genome proteins or Pfam domains. Its latest version features over 106,000 distinct protein families, each containing at least 100 sequences, cumulating to ∼20 million proteins.

As the number of protein sequences grows, the manual curation of protein families at scale seems unfeasible. Computational methods for automatically generating protein families must therefore balance sensitivity, scalability, and usability. Pfam-B [20], a compendium to the profile HMM based families in Pfam, is a computationally generated set of putative protein families that complement the Pfam families. Pfam-B families are based on a bidirectional MMseqs2 clustering [21], a very fast tool for sequence clustering, and are recalculated at each Pfam major release, but are not readily transferable to other sequences. Another widely used clustering tool is CD-HIT [22], which, similarly to MMseqs2, prioritizes speed by employing greedy incremental clustering starting from longer sequences. However, both MMseqs2 and CD-HIT sacrifice sensitivity compared to profile HMM methods, as well as present some challenges when wanting to transfer family annotations to additional sequences. Thus, there is a need for streamlined, scalable solutions that can generate and maintain protein families from large, and ever-expanding, sequence datasets.

We have developed the *nf-core/proteinfamilies* pipeline to address many of the aforementioned challenges by chaining modules for clustering sequences, generating protein family-level models, alignments and metadata, removing redundancies, and updating families as more sequences become available. The pipeline leverages the Nextflow [23] workflow orchestrator and nf-core principles [24,25], enabling standardized pipeline execution, efficient resource management, and parallel processing, hence allowing users to handle large datasets effectively.

## Methodology

The *nf-core/proteinfamilies* pipeline is a bioinformatics tool designed to generate new protein families or update existing ones, given a FASTA file of amino acid sequences as input, along with existing family HMMs and alignments when updating. The pipeline main components are *(i)* quality check, *(ii)* optional existing family model and alignment updating, *(iii)* sequence clustering, *(iv)* family generation, and *(v)* optional redundancy removal (Figure 1).

**Figure 1.**
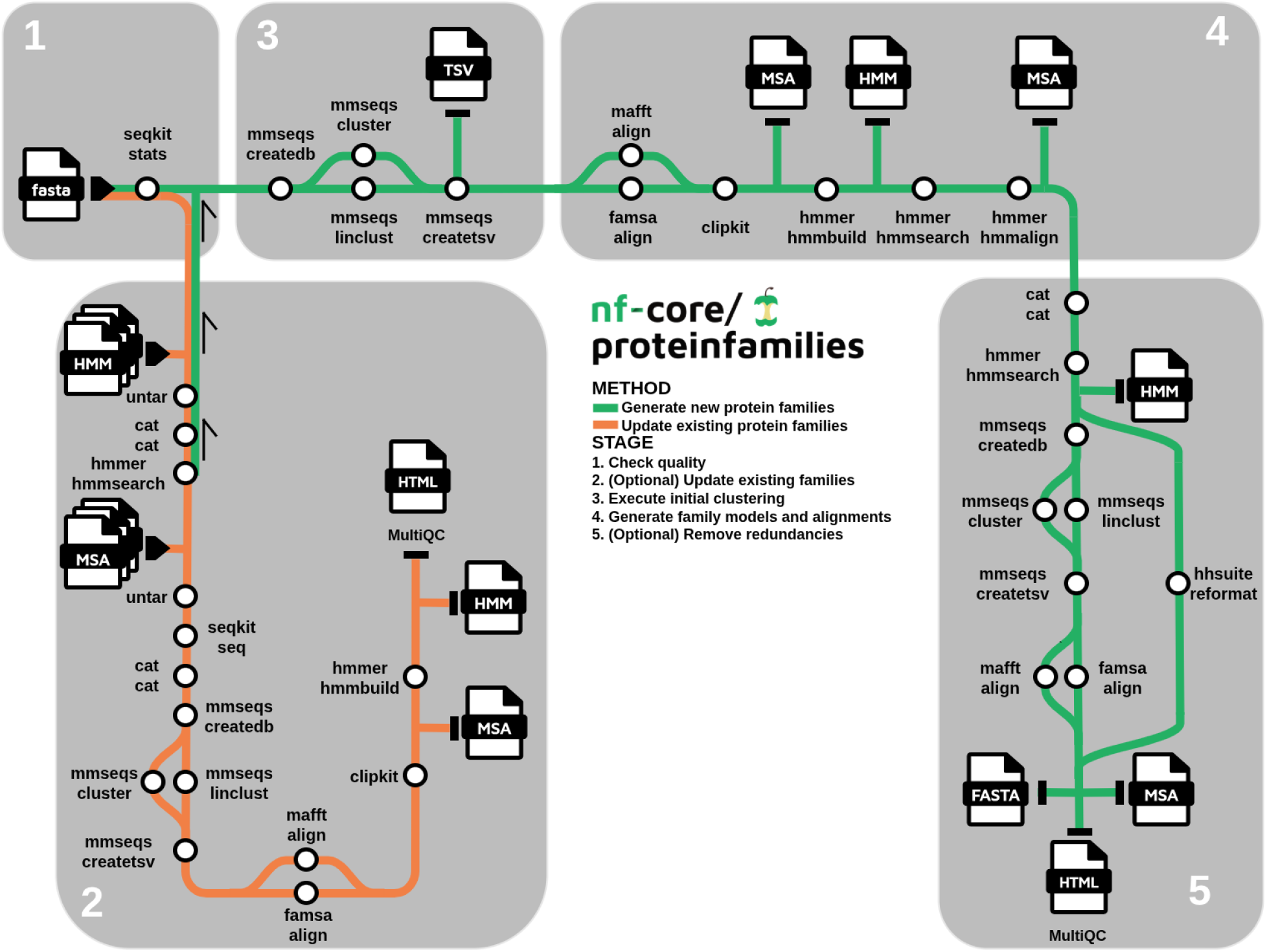
Workflow of the *nf-core/proteinfamilies* pipeline. The process begins with a mandatory input amino acid FASTA file, which undergoes a quality check using SeqKit (1) to generate sequence quality statistics. Optionally, in the *update existing families* stage (2), users can provide compressed archives containing existing family HMMs and MSAs. Sequences that match existing families are integrated into those families, updating their MSAs and HMMs accordingly. Sequences without matches follow the default workflow (green path) to generate new families, starting with an initial MMseqs2 clustering (3), followed by family generation steps (4). Finally, users may optionally apply redundancy reduction (5), either across families (inter-family) or within families (intra-family), to remove similar families or sequences, respectively.

### Quality check

The first stage employs a basic sequence quality check mechanism using SeqKit [26] which generates a report summarizing statistics such as the number of sequences, total amino acid count, and the minimum, average, and maximum sequence lengths.

### Existing families update

In this complementary workflow, users can enrich existing families with new sequences by recruiting new members into the full alignment of a family. When users provide existing family profile HMMs and multiple-sequence alignment (MSA) files*–*in compressed archive format (.tar.gz)*–*along with an amino acid sequence file, the pipeline begins by searching for matches of the input sequences against the current families. This is done by combining the existing HMMs into a library, and performing an *hmmsearch* against that with the input sequences. For each family hit, the relevant input sequences are combined with the non-aligned sequences from the corresponding family MSA in an aggregated FASTA file, which can optionally undergo strict clustering to eliminate redundancies. The remaining sequences are then aligned and optionally undergo gap removal, resulting in an updated family full MSA. Finally, the family profile HMM is retrained based on the updated MSA.

### Sequence clustering

The first stage of the workflow clusters protein sequences using MMseqs2, which enables fast sequence clustering. The input FASTA file is first converted into an MMseqs2 database, and then the MMseqs2 clustering algorithm of choice (*linclust* –focus on speed, *or cluster –*focus on sensitivity) is applied to group similar sequences into initial *seed* clusters. The clustering results are saved as a two-column (i.e., cluster representative, cluster member) TSV formatted output, and are then filtered based on minimum membership.

### Family generation

The next stage of the workflow generates protein families for each of the filtered clusters. In parallel, for each *seed* cluster, a multiple-sequence alignment is performed using either the *FAMSA* [27] or the *MAFFT* [28] aligner. This process generates the seed MSA file of a family. An optional step allows users to trim poorly aligned or gap-rich regions from the seed MSA using *ClipKIT* [29], either throughout the whole length of the alignment or only at the ends, thereby improving the alignment quality for downstream analyses. The seed alignments are then used to generate profile HMMs via the *HMMER* (v3.4) *hmmbuild* command [30]. The *hmmsearch* command is optionally used to recruit additional sequences from the input FASTA file into the generated families, according to user-defined thresholds such as *e*-value or minimum matching HMM length. Subsequently, recruited sequences are realigned to the family HMM with the *hmmalign* tool, generating a full MSA file for each family.

### Redundancy removal

In most cases, the initial MMseqs2 clustering will not assign all related sequences in a single cluster, producing different clusters that might be evolutionarily related. With the subsequent steps of the pipeline it is possible that the same sequence is assigned to multiple clusters leading to representation of the same protein family multiple times. Furthermore, in very large sequence datasets, the presence of identical or highly similar sequences within a family can unnecessarily consume storage space without contributing additional functional, evolutionary, or structural variability. To address these issues, the third stage of the pipeline employs two distinct mechanisms for redundancy detection and elimination. These mechanisms can be deployed both *(i) inter-family*, where redundancies between families are removed, and *(ii) intra-family*, where redundant sequences (those with very similar identity and alignment coverage) within a family are eliminated. Users can opt to utilize both mechanisms in combination, separately, or skip this step.

The inter-family redundancy removal process involves several steps. First, all generated profile HMMs are combined into a single file to form a profile HMM library. Next, family representative sequences are searched against the profile HMM library using *hmmsearch*. The premise is that if a family representative sequence matches another family’s HMM, with a strict matching length threshold (e.g., 90%), it suggests a significant similarity between the two families, allowing one to be labeled as redundant (currently removing the family with the fewest sequences).

In the intra-family redundancy removal mechanism, the pipeline begins with stringent MMseqs2 based clustering of members (*default parameters: 0*.*9 identity, 0*.*9 coverage length, bidirectional*), with only the cluster representatives retained. This step removes duplicate or nearly identical sequences, reducing the size of a family, while preserving a subset of diverse sequences that represent the protein family.

The workflow finishes with the generation of reports including the input sequence quality check, the size distribution of the initial MMseqs2 generated clusters and the produced family data and metadata using *MultiQC* [31]. The size distribution section reports the number of initial clusters for each observed size (number of proteins), while the produced family report contains information such as unique family identifiers, family sizes, and family representative sequences along with their length. These results are presented in an interactive HTML report, providing users with a concise overview of the generated data.

### Parameterization

The execution flow is guided by various user-defined parameters. For clustering, users can choose between two MMseqs2 clustering algorithms: *(i)* the standard *‘cluster’* method, suitable for medium-sized inputs (thousands to a few million sequences), or *(ii) ‘linclust’* (default), which offers faster but less sensitive clustering for larger sequence datasets (thousands of millions to a few billion sequences). Additional clustering parameters include sequence identity, coverage of the query/target, and coverage mode (unidirectional or bidirectional), which can be configured separately for initial input sequence clustering and redundancy removal (strict clustering within families). Users can also define the minimum cluster size required for the downstream family generation.

There are two available multiple sequence alignment (MSA) algorithms: *(i)* FAMSA and *(ii)* MAFFT. FAMSA is the default option due to its balance of time efficiency, memory usage, and general accuracy, while MAFFT can still outperform it in terms of alignment accuracy in certain edge cases [32]. Users may trim the resulting MSA using the ClipKit software, either throughout its length, or only at the ends of the alignment. MSA columns exceeding a user-defined gap percentage threshold can be clipped from within the alignment. To expand sequence families, users are encouraged to search the input sequences against the initial family HMMs generated by seed MSAs. This can be achieved by using *HMMER/hmmsearch*, where users can specify an e-value cutoff and minimum length threshold.

Lastly, users may enable redundancy removal after families are constructed. This process is controlled by two settings: one that eliminates redundancy between families and another that removes redundancy within families. Enabling both options, which is the default in the pipeline, is recommended to prevent family and sequence duplication while preserving sequence diversity. The amino acid sequence files (FASTA) that are generated throughout the pipeline can also be saved in the results folder by setting the relevant parameters.

### Protein Family Reproducibility Benchmark

To evaluate the applicability of the pipeline, we explored how well the automatically generated protein families produced by the pipeline matched expert curated protein families from established databases, which have undergone extensive curation, including multiple rounds of search and refinement (Figure 2). To do so, we collected protein sequences from 50 families (each containing at least 25 sequence members) from each of the following databases: NCBIFAM [33], PANTHER [34], Pfam [20], and HAMAP [36]. To explore diverse biological families and to prevent overlaps occurring between the different families, each family was sampled from different branches of the InterPro hierarchy tree. The 200 families matched a total of 159,605 non-redundant sequences. To this sequence dataset, we added 10,000 additional sequences from UniProt-SwissProt and verified the lack of significant similarity to the pooled family proteins using a *DIAMOND BLASTp* search [36] using default parameters. This resulted in a final input set of 169,605 unique protein sequences.

**Figure 2.**
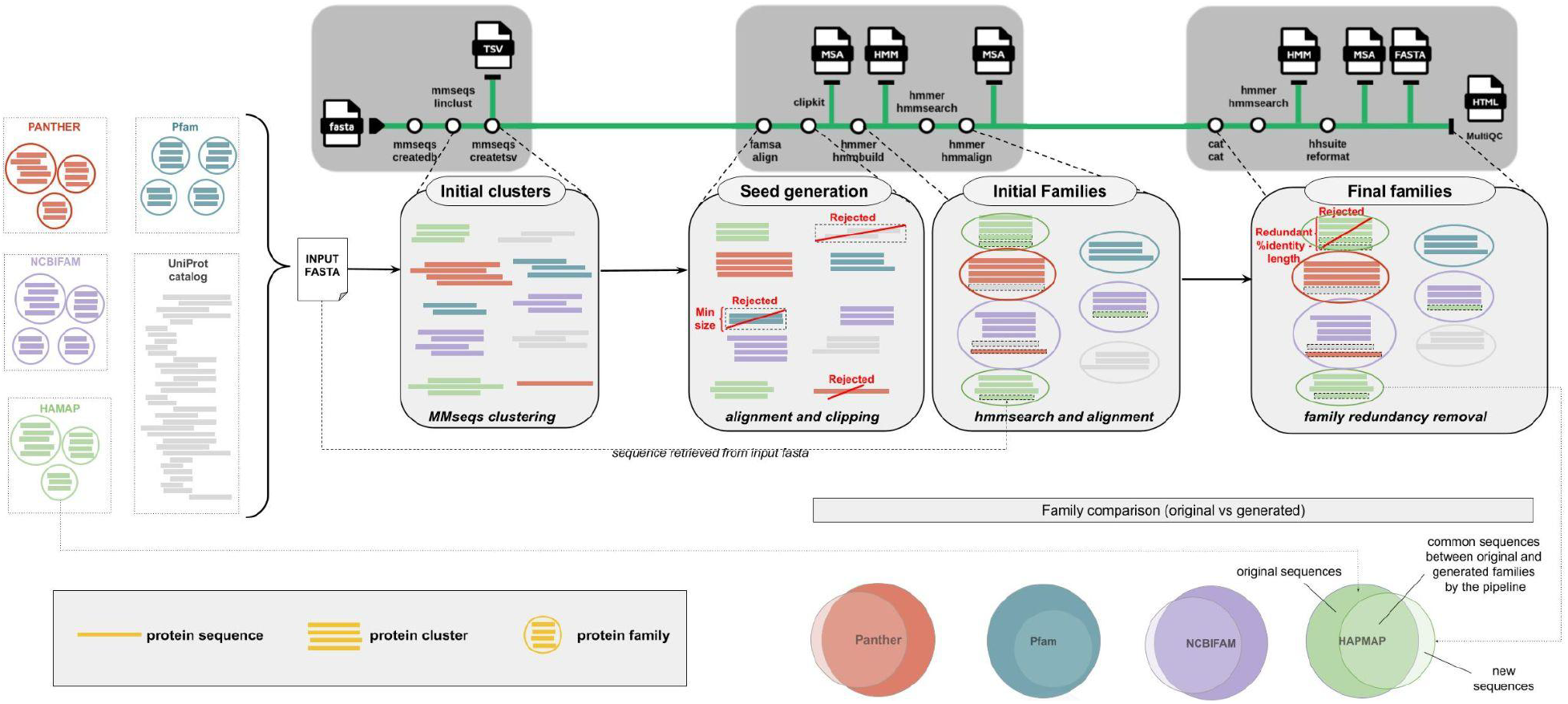
Workflow of the *nf-core/proteinfamilies* biological benchmark. First, 200 protein families are chosen from diverse InterPro entries and merged with a set of seemingly unrelated UniProt-SwissProt sequences as determined by a DIAMOND search. Next, the nf-core/proteinfamilies pipeline was applied to generate initial seed clusters with any cluster containing fewer than three sequences filtered out and MSA created - termed seed generation. Subsequently, the MSA is used to construct a profile HMM that is used to identify additional sequences related to the family. Finally, highly similar sequences are removed to reduce redundancy within the results. Overall, this process successfully incorporated 96.66% of the original family sequences into the output families, accurately reconstructing 173 of the 200 initial protein families.

Subsequently, we executed the nf-core/proteinfamilies pipeline with the combined protein dataset as input, using the ‘linclust’ algorithm along with a clustering identity of 0.5 and a minimum cluster size of 3 sequences, for the initial *seed* clusters generation. Using these parameters, the pipeline produced 2,947 *seed* clusters. The pipeline then recruited additional sequences into families via *hmmsearch*. Subsequently, families with a profile HMM match to another profile HMM across 100% of the length of the query HMM were eliminated to reduce redundancy, resulting in 709 remaining families. These 709 families captured 96.66% of the original unique sequence identifiers (103,385 out of 106,959). Even with this redundancy reduction step, a single input protein family can be represented by multiple families from the 709 families due to different length and match compositions. Thus, the 103,385 sequences were represented by 178,279 sequence regions. Family sequence recall was highest for HAMAP (99.7% of sequences, 84,051 / 84,335), followed by Pfam (90.6%, 7,452 / 8,227), NCBIFAM (89.9%, 2,407 / 2,677) and PANTHER (80.8%, 9,475 / 11,720). 2,941 additional UniProt sequences were also grouped into 295 families (without any sequence from the original input families set) out of the 709 (41.61%), and 395 out of the 709 (55.71%) contained only sequences from the original families. Despite the DIAMOND BLASTp search, 19 families contained both original input family and UniProt-SwissProt sequences.

To better understand how well these newly generated families represented the original, curated 200 families used as the input, we calculated the Jaccard index score between the protein identifiers contained in the overlap among produced and original families. 274 families out of the 414 new families containing input family sequences had a Jaccard index score of at least 0.5 (i.e., they contained at least 50% of the original family sequences), representing 173 distinct families out of the 200 original ones (Supplementary Table 1). Family representation across databases was highest for HAMAP (50/50), followed by NCBIFAM (44/50), Pfam (42/50), and PANTHER (37/50). 31 of the original families (15.5%) were represented by more than one new family, but differed in their sequence composition. An extreme example of such a case where an input family was represented by multiple families is PANTHER family PTHR23500, a sugar transport protein family, which is a long transmembrane family (average sequence length: 499) with 1,316 initial sequences. This input family was represented by 53 different families, each with slightly different sequence lengths due to the variability of the input *seed* alignments (but significantly overlapping, as they all pass the Jaccard score index).

27 of the original 200 families did not have any representation in the 274 families. 8 families were highly divergent, such that the MMseqs2 clustering could not produce seeds containing three or more sequences, hence were filtered out at that step. For example, NCBIFAM family TIGR01988 (IPR010971) had 31 original sequences, all of which were clustered as singletons, highlighting the diverse nature of the family. For another 16 families, while these were also divergent, initial *seed* alignments were produced, but were split into multiple small subfamilies such that the Jaccard similarity score with the original families was below 0.5 for each of them. For example, the PANTHER family PTHR23072 (IPR039527), was initially split into eight seed clusters with MMseqs2. After hmmsearch recruitment and family redundancy removal, two families remained; use_case_1124 and use_case_427, matching 19 and 31 sequences from PTHR23072 respectively, with no overlap between them, leaving 40 sequences unmatched. The Pfam family PF09826 (IPR019198) contains 68 total beta propellers and was split into use_case_4917 (30 sequences, 550 average length) and use_case_893 (16 sequences, 534 average length), with six sequences in common between the two generated families, leaving 28 initial sequences unmatched. The PANTHER family PTHR21227 (IPR006676), with an average sequence length of 347 amino acids, was mapped by three generated families; use_case_1769, use_case_2952 and use_case_2953. Families use_case_1769 and use_case_2953 were similar in length (244 and 239 respectively) and shared 52 of their total 54 respective sequences, whereas use_case_2952 recruited 20 of the larger original sequences, leading to an average sequence length of 459. Finally, three original families (PF00459, PF09704 and PTHR33910), were each partially mapped by a single generated family, but with a Jaccard Score below 0.5. The generated family use_case_118 matched 21 proteins from PF00459, leaving 22 unmatched, hence scoring just below the set threshold of 0.5. Similarly, use_case_3397 matched 14 out of 37 proteins from PF09704 leaving 23 unmatched, and use_case_2154 matched 8 out of 28 proteins from PTHR33910, leaving 20 unmatched.

Focusing on those families that scored above the Jaccard index score threshold, 255 families out of the 274 matching families contained only original family sequences (Supplementary Table 2). The remaining 19 families (2.68% of the total 709) contained a total of 27 sequences not found in the original families (note these are different to the 19 that fell below the Jaccard score index). Concerned that the families were recruiting sequences that were seemingly unrelated, we investigated them in more detail to understand the nature of the matches (Supplementary Table 3). The family use_case_3613, based on NCBIFAM NF045717, matched three sequences from PTHR23500 (A0A3B6G0N7/23-139, A0A3B6U448/23-139 and A0A3B6ETY9/23-140) and three further sequences from UniProt-SwissProt (O25918/1-120, T2KMF4/1116-1235 and Q1QI44/18-124). NF045717 is a family that contains response regulator receiver domain proteins without a DNA-binding domain, but all of the matched sequences also contain the same response regulatory domain. They have been excluded from this NCBIFAM presumably because the sequences in this family only have the Response Regulator domain, while the additional sequences contain additional accessory domains. The family use_case_2043, based on NCBIFAM TIGR03685, matched a sequence from PTHR20856 (A0A2G3AJI4/1-108) and another from UniProt-SwissProt (P10622/5-104). The TIGR03685 family contains large ribosomal subunit proteins from Archaea, and P10622 is also a large ribosomal subunit protein, explaining the recruitment. The PTHR20856 family contains longer sequences which are dna-directed RNA polymerase subunits, involved in RNA synthesis mechanisms. Nevertheless, the matched region of A0A2G3AJI4, residues 1-108, has a similar predicted structure to the sequences found in TIGR03685 sequences; a disorder region followed by four short helixes. The remaining 17 families contained either one or two sequences from the UniProt-SwissProt set. However, in all cases, the addition of these sequences to the family could be explained due to the presence of a common sequence motif. For example, use_case_1007 recruited all 2622 proteins from the HAMAP family of bacterial small ribosomal subunit proteins (uS14) (HAMAP accession: MF_01364_B), along with one additional UniProt-SwissProt sequence (A6MMU2). This sequence is also a small ribosomal subunit protein (uS14c) from the species *Illicium verum*, but neither the DIAMOND search nor the HAMAP entry identified this similarity.

### Computational Benchmark

Following the protein family reproducibility benchmark, we then evaluated the computational performance of the *nf-core/proteinfamilies* workflow. To do so, the pipeline (version 1.0.0) was tested with UniRef90, as a larger-scale benchmark, comprising 194,348,997 sequences (45GB compressed FASTA format). The workflow execution was conducted on a SLURM High-Performance Computing (HPC) infrastructure, utilizing Singularity containers. The shared HPC cluster comprises 340 compute nodes, each equipped with 48 physical cores and 468 GB of RAM, along with 20 memory nodes, each containing 48 physical cores and 1977 GB of RAM. One of the most resource-intensive stages of the pipeline is the initial MMseqs2 *linclust* clustering. A sequence identity of 50% and coverage of 90% for both strands were set as parameters. During this process, the pipeline utilized 150 GB of RAM. This step required the highest number of CPUs (12) within the pipeline and took 93.5 minutes to finish. Upon completion, clusters with less than 100 members were filtered out, since the larger ones are more likely to yield high-quality *seed* alignments. This step produced 23,245 seed clusters. After the inter-family redundancy removal (>80% model length similarity), 4,921 families remained (∼21% of the initial clusters). This reduction is expected, since we used a relaxed length threshold for model similarity (80%), that eliminated families which could have been sub-domains or super-families of other families, avoiding protein identifier duplication across the final set of families. Two histograms displaying the distribution of family sizes and representative sequence lengths are shown in Supplementary Figure 2.

Regarding the longest job durations, the FASTA partitioning process took approximately 3.3 hours, whereas the longest-running module was a FAMSA alignment process on an input containing over a million proteins, requiring 33 hours to complete. On average, each FAMSA process took 4.7 hours, each *hmmsearch* process required 83 minutes, and extracting family representative sequences, that can be used for further downstream analyses (i.e., protein annotation, structural prediction), from Stockholm MSA files took 37 hours. Detailed resource statistics can be found in the Supplementary File 1.

## Discussion

The ever-increasing size of protein databases limits the extent to which manual efforts for protein family curation can be applied. The open source *nf-core/proteinfamilies* pipeline provides essential features that make it well-suited for automated and scalable protein family generation. It also ensures portability, modularity, reproducibility, transparency and updating. The pipeline design prioritizes interoperability, producing standardized outputs at every stage to allow smooth integration with other tools and workflows. The pipeline also leverages Nextflow’s caching mechanism, which allows interrupted runs to resume without recomputing completed steps. Hosted by the nf-core organization on GitHub, the pipeline can be readily executed on any machine using Nextflow and Java. Users can choose to run it with Conda environments *(conda-forge/bioconda)* [37] or using containerized modules via Docker [38] or Singularity [39], enabling seamless deployment across different operating systems and computational environments. With comprehensive documentation and support for both the command-line interface (CLI) and the nf-core graphical user interface (GUI), researchers of various skill levels in bioinformatics can navigate, configure and use the pipeline effectively without extensive training.

The computational benchmarks (pipeline version 1.0.0) demonstrated that despite handling extremely large datasets, the resource requirements remained moderate, but did require access to a HPC. The resource utilisation statistics also highlighted key bottlenecks in the pipeline, such as the execution time required for extracting family representatives (required for post-pipeline downstream analyses). Subsequent to running this benchmark, we have addressed this performance issue, optimizing the module for extracting family representatives to enable it to be executed in parallel (as of v1.3.0). As this is a purely technical improvement that does not alter the output, we have not rerun our benchmarks with the updated version of the pipeline. While the pipeline currently scales well, further optimizations such as improved parallel execution, the use of more efficient sequence processing libraries, and dataset size-specific resource allocation, could further enhance computational performance, enabling continued scaling well into the future as sequence sets continue to grow.

As demonstrated by our protein family reproducibility benchmark, what constitutes a protein family differs according to the goal of the protein family model, for example capturing isofunctional sequences versus broad capture of a functional fold or full length sequences versus globular domains. Due to the nature of the initial seed generation, our approach will tend to focus on matching conserved regions or domains, rather than matching full length proteins due to the enabled trimming of alignments. Nevertheless, we were able to recreate a substantial set of protein families by placing 96.66% of sequences drawn from 200 InterPro families into 414 generated families, with only 19 of those recruiting additional UniProt-SwissProt sequences. Closer inspection of the 27 UniProt-SwissProt sequences matching the families revealed significant sequence similarity and/or consistent annotations indicating that these were not false positive matches. 27 of the original 200 families were not represented in the final set, as highly divergent families could not be created using the parameters in the benchmark, and are likely to be a limitation of the approach. Similarly, 31 original families were represented by more than one family, indicating that some of the curated families represented more divergent families, and that our unsupervised approach was unable to recapitulate them. Both types of families typically came from PANTHER or Pfam, which are recognised as representing more divergent families or domains. With the advent of protein structure based similarity searches sparked by the protein structure prediction revolution, these more distant relationships can now often be readily detected, overcoming this limitation.

The automatic generation of protein families at scale allows vast collections of sequences to be explored, as well as create starting points for manual curation efforts. Redundancy removal mechanisms can reduce storage requirements while retaining both family and sequence diversity, while the family updating mechanism enables updating of alignments and models when new sequences become available, without the need to recluster the entire sequence database and allow persistent identifiers to be established. With its robustness, scalability, portability, and ease of use, we anticipate that the *nf-core/proteinfamilies* pipeline will be widely adopted for protein family generation across various bioinformatics applications, further advancing research in the field.

## Supporting information

Supplementary Figure 1: Stacked barplots of reconstructed original families. Setting various Jaccard Scores as similarity thresholds, one can explore

Supplementary Figure 2: Histograms display the distribution of family sizes (capped at 8000) and representative sequence lengths, binned in increments

Supplementary File 1. Resource requirements for a large-scale computational benchmark with UniRef90 (194 million proteins). The resource diagrams illu

Supplementary Table 1: Jaccard index similarity scores among the protein sets of generated and original families. Only scores of at least 0.5 are repo

Supplementary Table 2. The generated families and their respective recruited additional UniProt-SwissProt sequences, sorted by descending additional U

Supplementary Table 3. The 19 generated families with mixed original family and additional sequence composition. The table shows the matched original

## Availability

The *nf-core/proteinfamilies* pipeline is implemented using Nextflow DSL2 and follows nf-core pipeline standards. The source code is available on GitHub at https://github.com/nf-core/proteinfamilies, and each release is archived on Zenodo (current latest release: 1.3.0, DOI: 10.5281/zenodo.14881993). For HPC infrastructures, the pipeline supports one of the available centralized nf-core configurations (currently more than 150): https://github.com/nf-core/configs. The pipeline can be executed either via CLI or through the nf-core GUI on the same website. Detailed documentation is available at https://nf-co.re/proteinfamilies. The computational benchmark results (MSA and HMM files) can be found at: https://zenodo.org/records/14997773. The reproducibility benchmark original family FASTA files, additional UniProt-SwissProt sequences, and generated family models and alignments can be found at: https://zenodo.org/records/16634647.

## References

1. Aplakidou E, Vergoulidis N, Chasapi M, Venetsianou NK, Kokoli M, Panagiotopoulou E, et al. Visualizing metagenomic and metatranscriptomic data: A comprehensive review. Comput Struct Biotechnol J. 2024; doi: 10.1016/j.csbj.2024.04.060.

2. Baltoumas FA, Karatzas E, Paez-Espino D, Venetsianou NK, Aplakidou E, Oulas A, et al. Exploring microbial functional biodiversity at the protein family level-From metagenomic sequence reads to annotated protein clusters. Front Bioinform. 2023; doi: 10.3389/fbinf.2023.1157956.

3. The UniProt Consortium, Bateman A, Martin M-J, Orchard S, Magrane M, Ahmad S, et al. UniProt: the Universal Protein Knowledgebase in 2023. Nucleic Acids Research. 2023; doi: 10.1093/nar/gkac1052.

4. Berman HM. The Protein Data Bank. Nucleic Acids Research. 2000; doi: 10.1093/nar/28.1.235.

5. O’Leary NA, Wright MW, Brister JR, Ciufo S, Haddad D, McVeigh R, et al. Reference sequence (RefSeq) database at NCBI: current status, taxonomic expansion, and functional annotation. Nucleic Acids Res. 2016; doi: 10.1093/nar/gkv1189.

6. Sayers EW, Cavanaugh M, Clark K, Pruitt KD, Schoch CL, Sherry ST, et al. GenBank. Nucleic Acids Research. 2022; doi: 10.1093/nar/gkab1135.

7. Chen I-MA, Chu K, Palaniappan K, Ratner A, Huang J, Huntemann M, et al. The IMG/M data management and analysis system v.7: content updates and new features. Nucleic Acids Research. 2023; doi: 10.1093/nar/gkac976.

8. Jumper J, Evans R, Pritzel A, Green T, Figurnov M, Ronneberger O, et al. Highly accurate protein structure prediction with AlphaFold. Nature. 2021; doi: 10.1038/s41586-021-03819-2.

9. Richardson L, Allen B, Baldi G, Beracochea M, Bileschi ML, Burdett T, et al. MGnify: the microbiome sequence data analysis resource in 2023. Nucleic Acids Research. 2023; doi: 10.1093/nar/gkac1080.

10. Mistry J, Chuguransky S, Williams L, Qureshi M, Salazar GA, Sonnhammer ELL, et al. Pfam: The protein families database in 2021. Nucleic Acids Res. 2021; doi: 10.1093/nar/gkaa913.

11. Paysan-Lafosse T, Blum M, Chuguransky S, Grego T, Pinto BL, Salazar GA, et al. InterPro in 2022. Nucleic Acids Res. 2023; doi: 10.1093/nar/gkac993.

12. Baltoumas FA, Karatzas E, Liu S, Ovchinnikov S, Sofianatos Y, Chen I-M, et al. NMPFamsDB: a database of novel protein families from microbial metagenomes and metatranscriptomes. Nucleic Acids Res. 2024; doi: 10.1093/nar/gkad800.

13. Pavlopoulos GA, Baltoumas FA, Liu S, Selvitopi O, Camargo AP, Nayfach S, et al. Unraveling the functional dark matter through global metagenomics. Nature. 2023; doi: 10.1038/s41586-023-06583-7.

14. Sillitoe I, Cuff AL, Dessailly BH, Dawson NL, Furnham N, Lee D, et al. New functional families (FunFams) in CATH to improve the mapping of conserved functional sites to 3D structures. Nucleic Acids Research. Oxford University Press (OUP); 2012; doi: 10.1093/nar/gks1211.

15. Huerta-Cepas J, Szklarczyk D, Heller D, Hernández-Plaza A, Forslund SK, Cook H, et al. eggNOG 5.0: a hierarchical, functionally and phylogenetically annotated orthology resource based on 5090 organisms and 2502 viruses. Nucleic Acids Research. 2019; doi: 10.1093/nar/gky1085.

16. Cantalapiedra CP, Hernández-Plaza A, Letunic I, Bork P, Huerta-Cepas J. eggNOG-mapper v2: Functional Annotation, Orthology Assignments, and Domain Prediction at the Metagenomic Scale. Tamura K, editor. Molecular Biology and Evolution. 2021; doi: 10.1093/molbev/msab293.

17. Kanehisa M, Sato Y, Kawashima M, Furumichi M, Tanabe M. KEGG as a reference resource for gene and protein annotation. Nucleic Acids Res. 2016; doi: 10.1093/nar/gkv1070.

18. Galperin MY, Vera Alvarez R, Karamycheva S, Makarova KS, Wolf YI, Landsman D, et al. COG database update 2024. Nucleic Acids Research. 2024; doi: 10.1093/nar/gkae983.

19. Tatusov RL, Fedorova ND, Jackson JD, Jacobs AR, Kiryutin B, Koonin EV, et al. The COG database: an updated version includes eukaryotes. BMC Bioinformatics. 2003; doi: 10.1186/1471-2105-4-41.

20. Paysan-Lafosse T, Andreeva A, Blum M, Chuguransky SR, Grego T, Pinto BL, et al. The Pfam protein families database: embracing AI/ML. Nucleic Acids Res. 2025; doi: 10.1093/nar/gkae997.

21. Steinegger M, Söding J. MMseqs2 enables sensitive protein sequence searching for the analysis of massive data sets. Nat Biotechnol. 2017; doi: 10.1038/nbt.3988.

22. Fu L, Niu B, Zhu Z, Wu S, Li W. CD-HIT: accelerated for clustering the next-generation sequencing data. Bioinformatics. Oxford University Press (OUP); 2012; doi: 10.1093/bioinformatics/bts565.

23. Di Tommaso P, Chatzou M, Floden EW, Barja PP, Palumbo E, Notredame C. Nextflow enables reproducible computational workflows. Nat Biotechnol. 2017; doi: 10.1038/nbt.3820.

24. Ewels PA, Peltzer A, Fillinger S, Patel H, Alneberg J, Wilm A, et al. The nf-core framework for community-curated bioinformatics pipelines. Nat Biotechnol. 2020; doi: 10.1038/s41587-020-0439-x.

25. Langer BE, Amaral A, Baudement M-O, Bonath F, Charles M, Chitneedi PK, et al. Empowering bioinformatics communities with Nextflow and nf-core.

26. Shen W, Sipos B, Zhao L. SeqKit2: A Swiss army knife for sequence and alignment processing. iMeta. Wiley; 2024; doi: 10.1002/imt2.191.

27. Deorowicz S, Debudaj-Grabysz A, Gudyś A. FAMSA: Fast and accurate multiple sequence alignment of huge protein families. Sci Rep. 2016; doi: 10.1038/srep33964.

28. Katoh K, Standley DM. MAFFT multiple sequence alignment software version 7: improvements in performance and usability. Mol Biol Evol. 2013; doi: 10.1093/molbev/mst010.

29. Steenwyk JL, Buida TJ, Li Y, Shen X-X, Rokas A. ClipKIT: A multiple sequence alignment trimming software for accurate phylogenomic inference. PLoS Biol. 2020; doi: 10.1371/journal.pbio.3001007.

30. Finn RD, Clements J, Eddy SR. HMMER web server: interactive sequence similarity searching. Nucleic Acids Res. 2011; doi: 10.1093/nar/gkr367.

31. Ewels P, Magnusson M, Lundin S, Käller M. MultiQC: summarize analysis results for multiple tools and samples in a single report. Bioinformatics. 2016; doi: 10.1093/bioinformatics/btw354.

32. Santus L, Garriga E, Deorowicz S, Gudyś A, Notredame C. Towards the accurate alignment of over a million protein sequences: Current state of the art. Curr Opin Struct Biol. 2023; doi: 10.1016/j.sbi.2023.102577.

33. Li W, O’Neill KR, Haft DH, DiCuccio M, Chetvernin V, Badretdin A, et al. RefSeq: expanding the Prokaryotic Genome Annotation Pipeline reach with protein family model curation. Nucleic Acids Res. 2021; doi: 10.1093/nar/gkaa1105.

34. Thomas PD, Kejariwal A, Campbell MJ, Mi H, Diemer K, Guo N, et al. PANTHER: a browsable database of gene products organized by biological function, using curated protein family and subfamily classification. Nucleic Acids Res. 2003; doi: 10.1093/nar/gkg115.

35. Lima T, Auchincloss AH, Coudert E, Keller G, Michoud K, Rivoire C, et al. HAMAP: a database of completely sequenced microbial proteome sets and manually curated microbial protein families in UniProtKB/Swiss-Prot. Nucleic Acids Res. 2009; doi: 10.1093/nar/gkn661.

36. Buchfink B, Xie C, Huson DH. Fast and sensitive protein alignment using DIAMOND. Nat Methods. 2015; doi: 10.1038/nmeth.3176.

37. Grüning B, Dale R, Sjödin A, Chapman BA, Rowe J, Tomkins-Tinch CH, et al. Bioconda: sustainable and comprehensive software distribution for the life sciences. Nat Methods. 2018; doi: 10.1038/s41592-018-0046-7.

38. Merkel D. Docker: lightweight Linux containers for consistent development and deployment. Linux J. 2014:2:22014;

39. Kurtzer GM, Sochat V, Bauer MW. Singularity: Scientific containers for mobility of compute. PLoS One. 2017; doi: 10.1371/journal.pone.0177459.

