## Supplementary figures and images for "nf-core/proteinfamilies: A scalable pipeline for the generation of protein families"

### Supplementary Figure 1: Stacked barplots of reconstructed original families. Setting various Jaccard Scores as similarity thresholds, one can explore

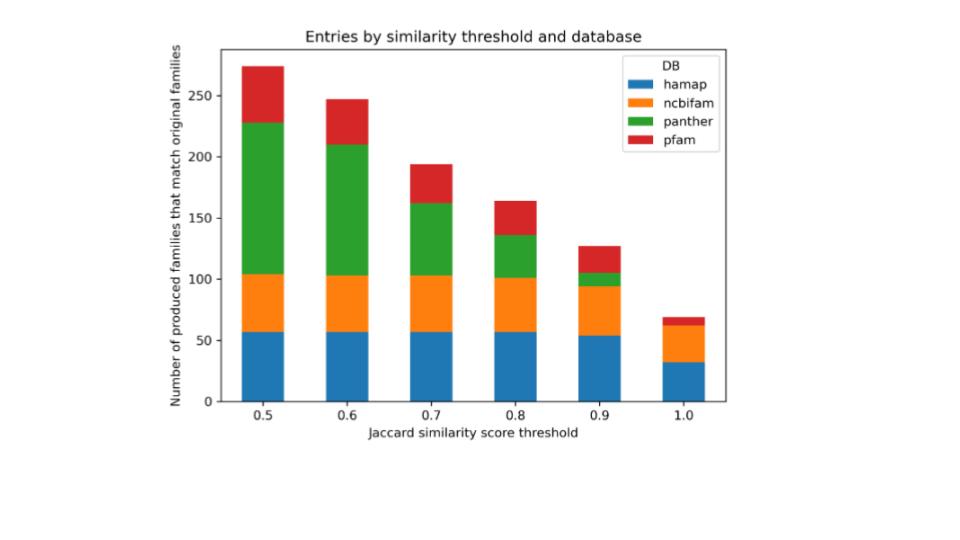

### Supplementary Figure 2: Histograms display the distribution of family sizes (capped at 8000) and representative sequence lengths, binned in increments

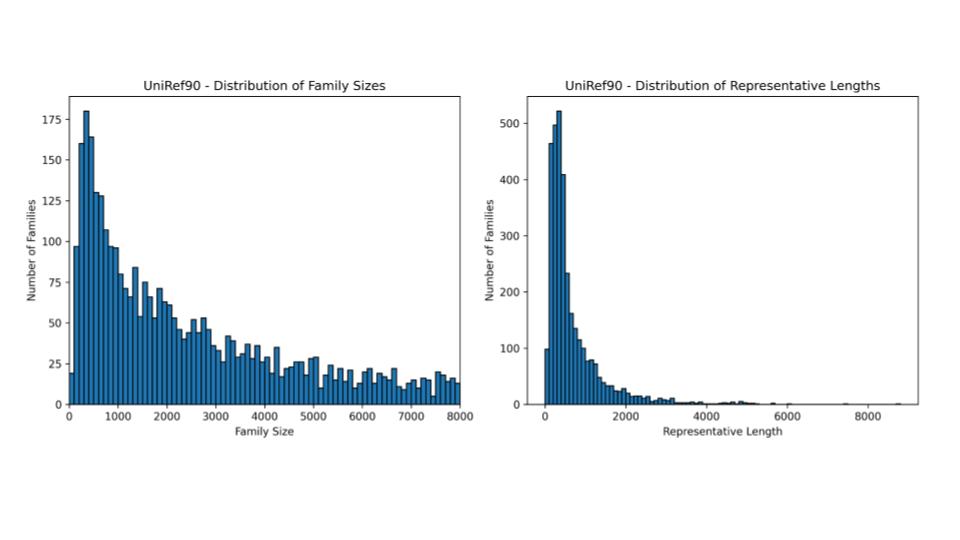

### Supplementary File 1. Resource requirements for a large-scale computational benchmark with UniRef90 (194 million proteins). The resource diagrams illu

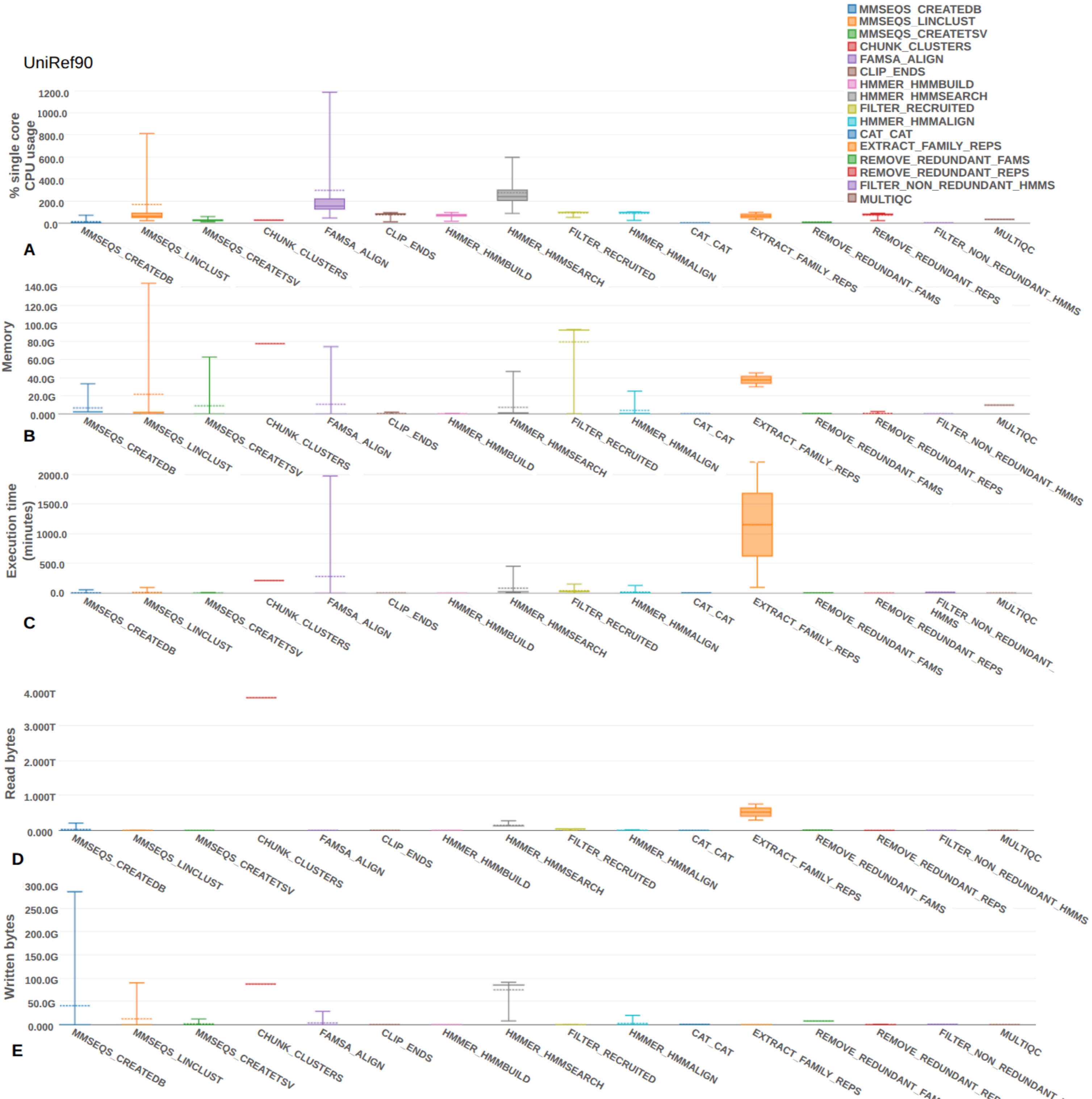
